# Top-down biological motion perception does not differ between adults scoring high versus low on autism traits

**DOI:** 10.1101/2023.09.08.556875

**Authors:** Danna Oomen, Jan R. Wiersema, Guido Orgs, Emiel Cracco

## Abstract

The perception of biological motion is an important social cognitive ability. Models of biological motion perception recognize two processes that contribute to the perception of biological motion: a bottom-up process that binds optic-flow patterns into a coherent percept of biological motion and a top-down process that binds sequences of body-posture ‘snapshots’ over time into a fluent percept of biological motion. The vast majority of studies on autism and biological motion perception have used point-light figure stimuli, which elicit biological motion perception predominantly via bottom-up processes. Here, we investigated whether autism is associated with deviances in the top-down processing of biological motion. For this, we tested a sample of adults scoring low *vs* high on autism traits on a recently validated EEG paradigm in which apparent biological motion is combined with frequency tagging (Cracco et al., 2022) to dissociate between two percepts: 1) the representation of individual body postures, and 2) their temporal integration into movements. In contrast to our hypothesis, we found no evidence for a diminished temporal body posture integration in the high-scoring group. We did, however, find a group difference that suggests that adults scoring high on autism traits have a visual processing style that focuses more on a single percept (i.e. either body postures or movements, contingent on saliency) compared to adults scoring low on autism traits who instead seemed to represent the two percepts included in the paradigm in a more balanced manner. Although unexpected, this finding aligns well with the autism literature on perceptual stability(/rigidity).

## Introduction

The ability to process human biological motion, and to extract intentions and affective states from it, plays a major role in social functioning (Pavlova, 2012). Hence, altered processing of biological motion may have serious social consequences. Based on this idea, an important hypothesis for why individuals with a diagnosis of autism spectrum disorder (henceforward ‘autism’) experience social difficulties (American Psychiatric Association, 2013) is that they process biological motion differently (Moore et al., 1997). Although individual studies investigating this hypothesis have so far yielded mixed results (group differences: e.g. Annaz et al., 2010; Koldewyn et al., 2010; Nackaerts et al., 2012; Price et al., 2012; no group differences: e.g. Cusack et al., 2015; Hubert et al., 2007; Saygin et al., 2010; Wright et al., 2014), three recent meta-analyses summarizing the literature all confirmed that biological motion perception is diminished in autism (Federici et al., 2020; Todorova et al., 2019; van der Hallen et al., 2019). However, the three meta-analyses also found high heterogeneity between studies, at least part of which is thought to be due to variations in the type of stimuli used to investigate biological motion perception (Federici et al., 2020; van der Hallen et al., 2019).

So far, most studies investigating biological motion perception in autism have used a class of stimuli referred to as ‘point-light figures’ (Federici et al., 2020). Point-light figures portray the movements of humans (or other animals) as a constellation of moving dots, typically placed on the major joints of the body (Johansson, 1973). These figures are popular because they are easy to adapt and convey kinematic information devoid of most form distractors (e.g. facial expression and looks). The processing of such point-light figures is known to rely on the integration of optic-flow patterns into a coherent percept of biological motion (Giese & Poggio, 2003; Johansson, 1973). This is referred to as bottom-up biological motion processing (or as the ‘motion’ pathway; Giese & Poggio, 2003; Lange & Lappe, 2006). Importantly, however, biological motion perception in real life also involves top-down biological motion processing (also referred to as the ‘form’ pathway), that is, the binding of body-posture ‘snapshots’ over time into a fluent movement percept (Giese & Poggio, 2003; Lange & Lappe, 2006). Because point-light figures minimize form information, they do not or only minimally engage this type of top-down biological motion processing (Blake & Shiffrar, 2007). In other words, while there is extensive research on bottom-up processing of biological motion in autism, we know only very little about top-down processing of biological motion in this group. This is surprising, because in real life, both types of processing contribute to how biological motion is perceived (Grossman & Blake, 2002) and differences in top-down processing could therefore play a role in the social difficulties associated with autism.

To directly investigate whether top-down processing of biological motion differs in autism, biological motion perception should ideally be studied with an *apparent* biological motion paradigm. In short, these paradigms represent movements as a sequence of purely static body-postures (Chatterjee et al., 1996; Shiffrar & Freyd, 1990). If these sequences are presented with a time interval between consecutive body postures that is biomechanically possible, this is known to elicit a percept of biological motion in the absence of retinal motion (Orgs & Haggard, 2011). Importantly, given that there is no retinal motion, these paradigms rely predominantly on top-down instead of bottom-up processing (e.g., Orgs et al., 2016). More specifically, it is assumed that apparent biological motion perception is the result of integrating static body postures over time (Giese & Poggio, 2003; Lange & Lappe, 2006).

Recently, a new electroencephalogram (EEG) paradigm was developed to capture this temporal binding process (Cracco et al., 2022). In the paradigm, apparent biological motion (Orgs et al., 2011, 2013) is combined with frequency tagging (Norcia et al., 2015) by showing sequences of 12 body postures (Figure 1), either in their natural order (fluent condition) or in a non-fluent order (non-fluent condition). Importantly, the sequences are symmetrical at the midpoint, which results in three frequencies of interest: one coupled to image presentation (base rate), one coupled to the symmetrical turning point in the sequence (half cycle; every 6th image), and one coupled to the repetition of the sequence (full cycle; every 12th image). Critically, fluent and non-fluent sequences generate a different primary percept. In fluent sequences, the primary percept is a series of cyclical movements centred around the half cycle point. Instead, in non-fluent sequences, it is a series of body postures centred around the full cycle point. Stated differently, movements repeat at half cycle rate and body posture sequences repeat at full cycle rate. As a result, their processing can be dissociated at different frequencies of the brain response: the representation of individual body postures is captured at full cycle frequencies, whereas their temporal integration into movements is captured at half cycle frequencies.

**Figure 1.**
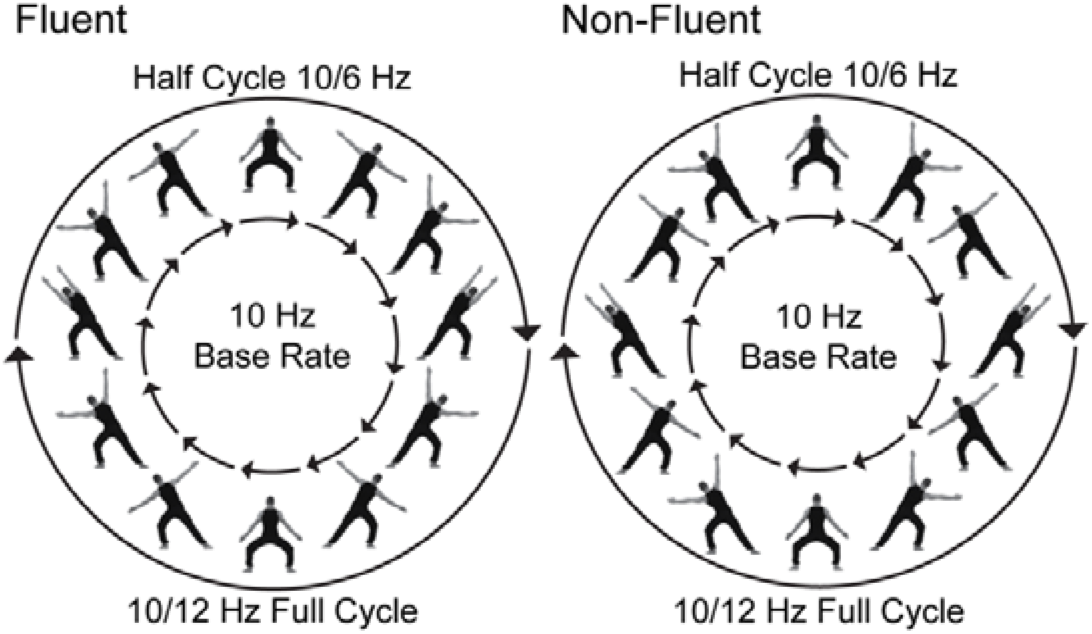
Stimulus sequence for the fluent and non-fluent condition. Images are presented at a base rate of 10 Hz. Individual postures are repeated at a frequency of 10/12 Hz (full cycle frequency), and movements are completed at a frequency of 10/6 (half cycle frequency). Figure adapted from Cracco et al. (2022; published under a CC BY license).

With this paradigm, Cracco et al. (2022) indeed found that brain responses at half cycle frequencies were stronger when the postures were ordered to form a fluent movement (fluent > non-fluent condition), whereas brain responses at full cycle frequencies were stronger when they were not (non-fluent > fluent condition; see also Cracco et al., 2023). Here, we applied the apparent biological motion task of Cracco et al. (2022) to investigate if temporal integration of body postures into movements differs between individuals scoring high *vs* low on autism traits. Autism traits are continuously distributed in the population (Abu-Akel et al., 2019) and previous research has shown that such a dimensional approach in the neurotypical population can give valuable insights into autism and its traits (e.g. Nijhof et al., 2017). If top-down biological motion perception is diminished in autism, one would expect the high-scoring group to show diminished temporal integration of body postures into movements. In other words, to show a reduced effect of movement fluency on half cycle responses but not on full cycle responses.

## Methods

### Participants

All participants had normal or corrected-to-normal vision, were proficient in Dutch, reported no known neurological condition, had not sought professional help for a mental health problem in the last 6 months before participation, and scored low (≤ 2) or high (≥ 6) on the autism-spectrum quotient-10 (AQ-10; Allison et al., 2012). The AQ-10 is a brief screening tool for ASD that contains 10 items (i.e. the two items with the greatest discriminatory power in each of the five subscales of the full AQ) with a recommended cut-off of 6 for clinical screening purposes. Participants were recruited via an online pre-screening questionnaire in which we assessed the inclusion criteria. A total of 1171 persons filled out the questionnaire, of whom 477 scored low and 84 scored high on the AQ-10. Of those people, 42 eligible candidates of each group (i.e., low- and high-scoring) agreed to participate. Three participants were excluded due to technical issues, and 10 participants due to bad signal quality^1^ (> 8 electrodes requiring interpolation), resulting in a final sample of 69 participants (34 low-scoring, 35 high-scoring participants). Groups did not differ in age, sex, or years of education (See Table 1). The experimental protocol was approved by the local ethics committee of the Faculty of Psychology and Educational Sciences of Ghent University (EC/2019/32), and informed consents were obtained from all participants prior to the study. The participants were compensated for their time.

**Table 1.** Participant characteristics.

|  | Low-scoring group<br>*M (SD)* | High-scoring group<br>*M (SD)* | *F*-value (*p*) |
| --- | --- | --- | --- |
| Age | 22.65 (3.59) | 21.46 (2.78) | 2.38 (.128) |
| Sex (# of females) | 17 | 17 | n/a |
| Years of education | 15.82 (1.85) | 15.57 (2.61) | 0.87 (.354) |
| Full AQ | 11.62 (4.2) | 22.06 (7.82) | 47.34 ($< .001$) |
| SRS-A | 30.29 (12.36) | 60.77 (25.24) | 40.19 ($< .001$) |

*Note.* low-scoring group $n = 34$ ; high-scoring group: $n = 35$ ; full AQ = autism-spectrum quotient; SRS-A = social responsiveness scale - adults
<sup>1</sup> The late identification of faulty equipment led to this high number of participants with bad signal quality.

### Task and Procedure

Participants were seated in an electrically shielded room approximately 60 cm from a 24-inch computer monitor. Before the start of the apparent biological motion task, all participants completed two questionnaires as well as a computer task of 16 minutes This task addressed an unrelated research question and is therefore not further reported here. After a break, participants completed the apparent biological motion task of approximately 22 minutes. This task was presented using PsychoPy3 (Peirce et al., 2019).

The AQ (full version; Baron-Cohen et al., 2001; Hoekstra et al., 2008) and the social responsiveness scale for adults (SRS-A; Constantino, 2002; Noens et al., 2012), two commonly used questionnaires to measure ASD traits, were administered at the start of the test sessions to further describe our sample. As expected, both the AQ and SRS-A scores differed significantly between groups (see Table 1) and correlated with the AQ-10 score (AQ: *r_s_*= .71, *p* < .001; SRS-A: *r_s_* = .70, *p* < .001).

The apparent biological motion task was nearly identical to the one described by Cracco et al. (2022). Participants were presented with repeating sequences of 12 images (12° × 12°) on a grey background, depicting a dancer in 12 successive whole-body postures. In the fluent condition, the body postures were ordered to produce a fluent movement percept. In the non-fluent condition, the same images were reordered to produce maximal visual displacement between body postures (see Figure 1). For both conditions, sequences were symmetrical at the midpoint, leading to three frequency responses coupled to distinct features of stimulus presentation: one coupled to image presentation of 10 Hz (base rate), one coupled to the symmetrical turning point in the sequence (half cycle., every 6^th^ image: 10 Hz/6 = 1.67 Hz), and one coupled to the repetition of the image sequence (full cycle, every 12^th^ image: 10 Hz/12 = 0.83 Hz). Crucially, in the fluent condition, the half cycle point is linked to movement completion from side-to-side and movement processing should therefore result in stronger half cycle responses. In contrast, the full cycle point is coupled to the completion of the full body sequence and processing body postures should therefore result in stronger full cycle responses instead.

The two conditions were presented 5 times in randomized blocks. A block consisted of a 120 s video with a 10 s fade in (0-100%) and a 10 s fade out (100-0%) to avoid abrupt eye movements and blinks due to the sudden (dis)appearance of stimuli. The videos were created by repeating the 12-image sequence 100 times. To stimulate attention, participants were instructed to fixate on a grey cross positioned in the centre of the screen, and to press the space bar every time it briefly (400 ms) turned red. The task started with a practice block, for which we used the random condition of Cracco et al. (2022), in which the body postures were presented at random. This control condition only elicits a base rate response as neither postures nor movements repeat predictably when images are presented at random. As Cracco et al. (2022) already demonstrated this, we did not include the random condition in the actual task. However, inspection of the practice block data confirmed that the control condition did indeed only elicit a response at base rate (See Supplementary Material).

### EEG Recording and Pre-processing

EEG was continuously recorded from 64 scalp sites using a sampling rate of 1000 Hz, an ActiCHamp amplifier (Brain Products, Enschede, The Netherlands), and BrainVisionRecorder software (version 1.21.0304, Brain Products, Gilching, Germany). Ag/AgCI active electrodes were positioned according to the extended 10-20 international system. During EEG recording, all channels were referenced to Fz. Horizontal EOG was recorded with FT9 and FT10 electrodes embedded in the cap. Vertical electro-oculogram (EOG) was recorded with additional bipolar AG/AgCI sintered ring electrodes placed above and below the left eye.

Off-line processing of the EEG signal was done using Letswave 6 (https://www.letswave.org/) and followed the procedure of Cracco et al. (2022). First, we applied a fourth-order Butterworth band-pass filter (0.1 Hz - 100 Hz), after which we segmented the data to obtain epochs extending from 2 s before to 122 s after the stimulus onset. Next, ocular artefacts were removed with an independent component analysis (ICA) on the merged segmented data using the Runica algorithm and a square matrix. For each participant, ICs were inspected visually and the ICs related to eye blinks were removed manually. After ICA, we interpolated noisy or faulty electrodes using data from the three closest neighbouring electrodes. The signal was then re-referenced to an average reference. After re-referencing, the segments were cropped to cut out the fade in and fade out periods. This resulted in segments of 96 s and caused the duration of a single segment to be a multiple of the duration of a single sequence (1.2 s), thereby ensuring that the frequencies of interest were captured by a single frequency bin. Finally, trials were averaged per condition and subsequently a Fast Fourier Transform (FFT) was applied to transform the data of each electrode to normalized (divided by N/2) amplitudes in the frequency domain.

### Analysis

For the statistical analysis, we computed the signal to noise-subtracted amplitudes (SNS) at each frequency bin by subtracting the average voltage amplitude of the 20 neighbouring bins as baseline (10 on each side, excluding the immediate adjacent bin). This was done for the three neural responses separately (i.e., base rate, full cycle, and half cycle). Based on the study by Cracco et al. (2022), the SNS for the base rate response was calculated as the sum of the harmonics at 10 Hz, 20 Hz, 30 Hz, 40 Hz, 60 Hz, 70 Hz, 80 Hz, and 90 Hz, the full cycle response was calculated as the sum of the harmonics at 0.83 Hz, 2.50 Hz, 4.17 Hz, 5.83 Hz, 7.50 Hz, 9.17 Hz, 10.83 Hz, 12.5 Hz, 14.17 Hz, and 15.83 Hz, and the half cycle response was calculated as the sum of the harmonics at 1.67 Hz, 3.33 Hz, 5.00 Hz, 6.67 Hz, 8.33 Hz, 11.67 Hz, 13.33 Hz, 15.00 Hz, 16.67 Hz, and 18.33 Hz. Note that only the odd harmonics of the full cycle response were included to avoid overlap with the half cycle response, that 10 Hz was excluded for the half cycle response to avoid overlap with the base rate response, and that 50 Hz and 100 Hz were excluded in the base rate response to exclude line noise.

Based on the study by Cracco et al. (2022), we included the SNS data of 4 clusters in our statistical analyses: a left posterior (PO3, PO7 O1), middle posterior (Poz, Oz), right posterior (PO4, PO8, O2), and a frontocentral cluster (FC1, FCz, FC2). Note that the electrodes included in each cluster were identical to the ones used by Cracco et al. (2022), with the exception that we did not include Iz in the middle posterior cluster, because our electrode setup did not record this electrode site.

On the SNS data, we performed separate mixed design ANOVAs for the base rate (10 Hz), full cycle (10/12 Hz), and half cycle (10/6 Hz) responses, using Condition (fluent, non-fluent) and Region (left posterior, middle posterior, right posterior, or middle central) as within-subject factors, and Group (low, high) as between-subject factor. Whenever violations of sphericity occurred, the ANOVA degrees of freedom were adjusted according to the Greenhouse-Geisser formula. Significant interactions and main effects of region were followed up by two-tailed *t*-tests. T-values are reported as absolute values.

## Results

### Base Rate (10 Hz)

Images are presented at the base rate of 10 Hz. The base rate therefore captures the processing of images or low-level visual processes associated with the image transitions (e.g., contrast change; Cracco et al., 2022, 2023). As the visual change from image to image is strongest in the non-fluent condition, the base rate response should be strongest for the non-fluent condition. The base rate analysis revealed a main effect of Condition, *F*(1, 67) = 10.36, *p* = .002, ηp² = .13, a main effect of Region, *F*(2.51, 168.14) = 120.07, *p* < .001, ηp² = .64, and a Condition × Region interaction, *F*(2.69, 180.46) = 6.21, *p* = .001, ηp² = .09. There was no main effect of Group, nor interaction effects with Group, all *p*s ≥ .292.

As expected, the main effect of Condition indicated that the base rate response was stronger in the non-fluent condition (*M* = 2.06, *SD* = 0.87) than in the fluent condition (*M* = 1.89, *SD* = 0.85), *d_z_* = 0.39, 95% CI [0.14, 0.63]. The main effect of Region revealed that the base rate response was stronger in the three posterior regions (left *M* = 2.11, *SD* = 0.98; middle *M* = 2.51, *SD* = 1.26; right *M* = 2.59, *SD* = 1.19) than in the central region (*M* = 0.71; *SD* = 0.36), all *t*(68) ≥ 15.03, *p* < .001, *d_z_* ≥ 1.81, and that the base rate response was stronger in the middle and right posterior region than in the left posterior region, both *t*(68) ≥ 3.58, *p* ≤ .001. However, no difference was found between the middle posterior region and the right posterior region *t*(68) = 0.63, *p* = .531, *d_z_* = 0.07, 95% CI [-0.16, 0.31].

The Condition × Region interaction showed that the effect of fluency was present in the left and right posterior regions (non-fluent left: *M* = 2.21, *SD* = 1.05; fluent left: *M* = 2.01, *SD* = 0.99; non-fluent right: *M* = 2.75, *SD* = 1.27; fluent right: *M* = 2.43,*SD*= 1.20), both *t*(68) ≥ 3.18, *p* ≤ .002, *d_z_* ≥ 0.38, and to a lesser extent in the central region (non-fluent: *M* = 0.73, *SD*= 0.39; fluent: *M* = 0.68, *SD*= 0.37), *t*(68) ≥ 2.00, *p* = .050, *d_z_* = 0.24, 95% CI [3.74×10^4^, 0.48], but not in the middle posterior region, *t*(68) = 1.18, *p* = .244, *d_z_* = 0.14, 95% CI [-0.10, 0.38].

### Full Cycle Rate (10/12 Hz)

Individual postures are repeated at the full cycle rate. The full cycle rate therefore primarily captures body perception. As the primary percept in the non-fluent condition is a repeating posture sequence, the full cycle response should be strongest for the non-fluent condition (Cracco et al., 2022, 2023). The full cycle analysis revealed a main effect of Condition *F*(1, 67) = 52.86, *p* < .001, ηp² = .44, a main effect of Region, *F*(2.65, 177.74) = 181.57, *p* < .001, ηp² = .73, a Region × Condition effect, *F*(2.46, 164.58) = 16.93, *p* < .001, ηp² = .20, and a Condition × Group effect, *F*(1, 67) = 4.27, *p* = .043, ηp² = .06. There was no main effect of Group or any other interaction effects, all *p*s ≥ .547.

As expected, the main effect of Condition indicated that the full cycle response was stronger in the non-fluent (*M* = 1.67, *SD* = 0.62) than in the fluent condition (*M* = 1.29, *SD* = 0.50), *d_z_* = 0.86, 95% CI [0.85, 1.13]. The main effect of Region showed that the response was stronger in the two lateral posterior regions (left *M* = 2.08, *SD* = 0.88; right *M* = 2.20,*SD* = 0.88) than in the middle posterior region (*M* = 1.33, *SD* = 0.63), both *t*(68) ≥ 8.76, *p* ≤ .001, *d_z_* ≥ 1.05, and stronger in the three posterior regions than in the central region (*M* = 0.30, *SD* = 0.14), all *t*(68) ≥ 14.65, *p* ≤ .001 *d_z_* ≥ 1.76. However, no difference was found between the left and right posterior regions, *t*(68) = 1.20, *p* = .236, *d_z_* = 0.14, 95% CI [-0.09, 0.38].

The Condition × Region interaction showed a stronger response in the non-fluent than in the fluent condition at every region, all *t*(68) ≥ 2.32, *p* ≤ .023, *d_z_* ≥ 0.28 (for means and standard deviations see Supplementary Table 1). The fluency effect was, however, stronger in the posterior regions than the central region, all *t*(68) ≥ 3.12, *p* ≤ .003, *d_z_* ≥ 0.38, and stronger in the lateral than in the middle posterior regions, both *t*(68) ≥ 3.23, *p* ≤ .002, *d_z_* ≥ 0.39. However, no difference between the two lateral posterior regions was found, *t*(68) = 0.42, *p* = .679, *d_z_* = 0.05, 95% CI [-0.19, 0.29].

The Condition × Group interaction showed that even though the full cycle response was stronger in the non-fluent condition (low: *M* = 1.62, *SD* = 0.61; high: *M* = 1.71, *SD* = 0.64) than in the fluent condition (low: *M* = 1.35, *SD* = 0.57; high: *M* = 1.23, *SD* = 0.43) for both groups, both *t*(≥ 33) ≥ 4.04, *p* ≤ .001 *d_z_* ≥ 0.69, this fluency effect was larger in the high-scoring group (*M* = 0.48, *SD* = 0.47) than in the low-scoring group (*M* = 0.27, *SD* = 0.39).

### Half Cycle Rate (10/6 Hz)

Movements are completed at the half cycle rate. The half cycle rate therefore primarily captures movement perception. As the primary percept in the fluent condition is a series of movements, the half cycle response should be strongest for the fluent condition (Cracco et al., 2022, 2023). The half cycle analysis revealed a main effect of Condition, *F*(1, 67) = 132.89, *p* < .001, ηp² = .67, a main effect of Region, *F*(2.34, 157.06) = 196.61, *p* < .001, ηp² = .75, a Condition × Region effect, *F*(1, 201) = 21.17, *p* < .001, ηp² = .24, a Condition × Group effect, *F*(1, 67) = 5.00, *p* = .029, ηp² = .07 and to a lesser extent a Condition × Region × Group effect, *F*(3, 67) = 2.83, *p* = .051, ηp² = .04. There was no main effect of Group or a Region × Group effect, both *p*s ≥ .662.

As expected, the main effect of Condition indicated that the half cycle response was stronger in the fluent condition (*M* = 2.86, *SD* = 1.02) than in the non-fluent condition (*M* = 1.90, *SD* = 0.76), *d_z_* = 1.35, 95% CI [-1.68, -1.02]. The main effect of Region showed that the half cycle response was stronger in the three posterior regions (left: *M* = 2.63, *SD* = 1.01; middle: *M* = 2.99, *SD* = 1.03; right: *M* = 3.03, *SD* = 1.27) than in the central region (*M* = 0.86, *SD* = 0.36), all *t*(68) ≥ 17.87, ≤ .001, *d_z_* ≥ 2.15, and that the response was stronger in the left posterior region than in the middle and right posterior regions, both *t*(68) ≥ 3.23, *p* ≤ .002, *d_z_* ≥ 0.39. However, no difference was found between the middle posterior region and the right posterior region, *t*(68) = 0.36, *p* = .719, *d_z_* = 0.04, 95% CI [-0.19, 0.28].

The Condition × Region interaction showed a fluency effect in every region, all *t*(68) ≥ 8.92, *p* ≤ .001, *d_z_* ≥ 1.07 (for means and standard deviations see Supplementary Table 2). The effect was, however, stronger in the posterior regions than in the central region, all *t*(68) ≥ 5.58, *p* < .001 *d_z_* ≥ 0.67, and slightly stronger in the right posterior region than in the middle posterior region, *t*(68) = 2.01, *p* = .049, *d_z_* ≥ 0.24, 95 CI [-0.48, 0.00]. No other region differences were found, both *p*s ≥ .164.

The Condition × Group interaction revealed that even though the half cycle response was stronger in the fluent condition (low: *M* = 2.79, *SD* = 0.97; high: *M* = 2.92, *SD* = 1.07) than in the non-fluent condition (low: *M* = 2.02, *SD* = 0.77; high: *M* = 1.78, *SD* = 0.73) for both groups, both *t* ≥ 6.66, *p* < .001, *d_z_* ≥ 1.14, this fluency effect was larger in the high-scoring group (*M* = 1.14, *SD* = 0.70) than in the low-scoring group (*M* = 0.77, *SD* = 0.68).

Finally, the Region × Condition × Group indicated that this larger fluency effect for the high *vs* the low-scoring group was restricted to the middle and right posterior regions, both *t*(67) ≥ 2.19, *p* < .032, *d* = 0.53 (for means and standard deviations see Supplementary Table 3).

**Figure 1.**
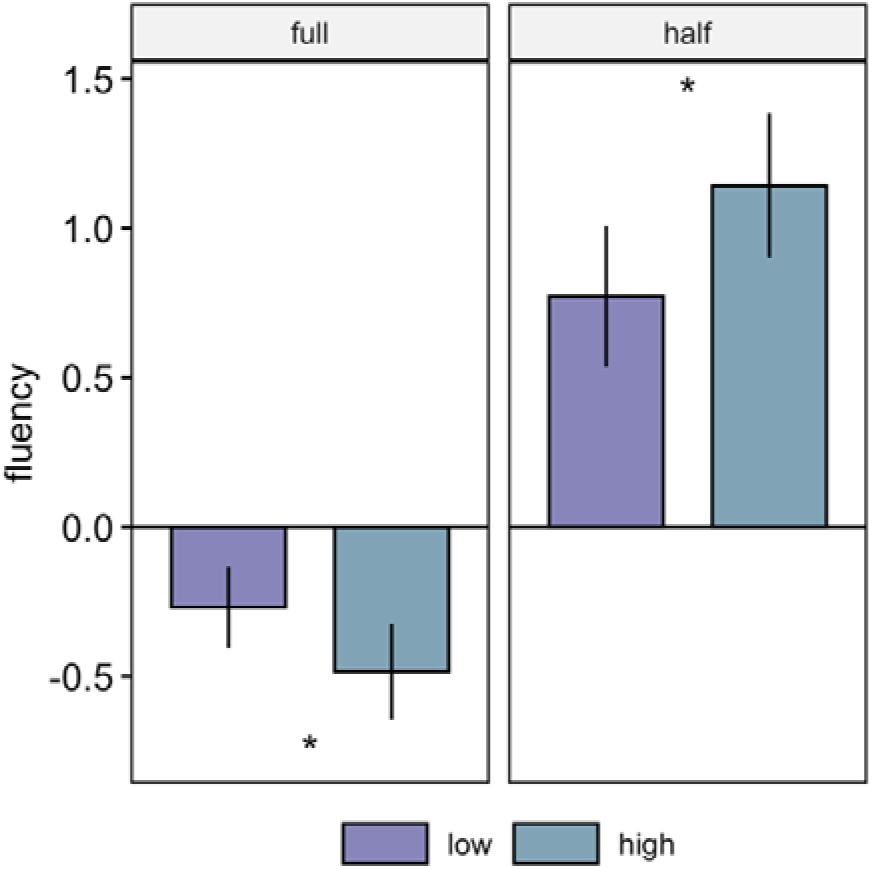
Fluency effects (fluent – non-fluent) for the two groups (low-scoring, high-scoring) and the two cycle responses (full, half) separately. Error bars represent between-subject 95% CI’s.

**Figure 2.**
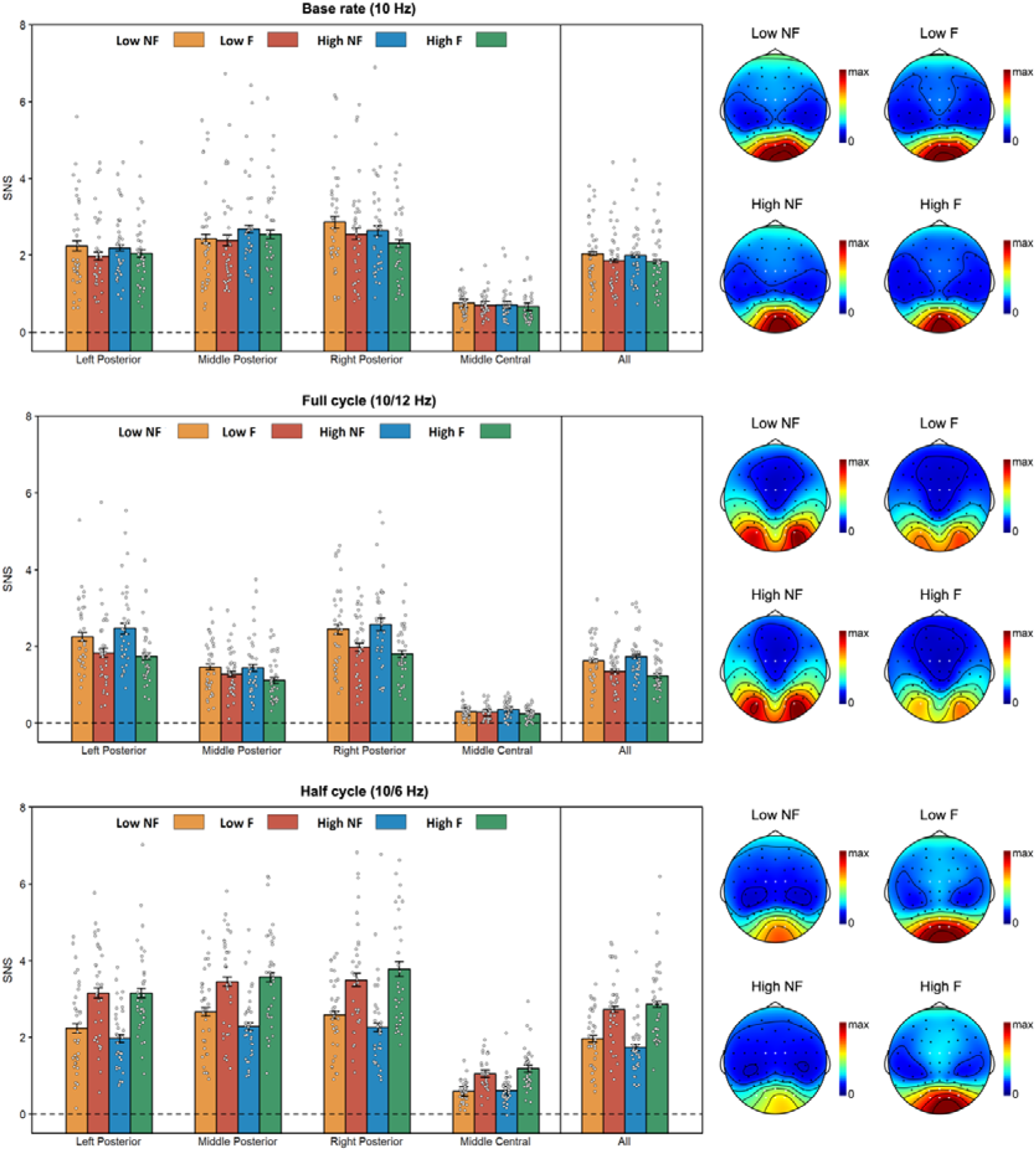
Noise-subtracted amplitudes (SNS) per group for the two conditions and separately for each region cluster, together with their topographies; Low = low-scoring group, High = high-scoring group, NF = non-fluent, F = fluent. Error bars are standard errors of the mean corrected for within-subject design according to Morey (2008). Topographies are scaled from 0 to the maximum amplitude across conditions for the respective response.

## Discussion

The perception of biological motion is an important social-cognitive ability. It has been hypothesised that autism is associated with altered biological motion perception, and that this difference contributes to social difficulties (Kaiser & Pelphrey, 2012; Moore et al., 1997; Pavlova, 2012). In line with this hypothesis, three independent meta-analyses recently confirmed that biological motion perception in autism is diminished (Federici et al., 2020; Todorova et al., 2019; van der Hallen et al., 2019). Two processes are involved in the perception of real-life biological motion (Grossman & Blake, 2002): a bottom-up process that binds optic-flow patterns into a coherent percept of biological motion and a top-down process that binds sequences of body-posture ‘snapshots’ over time into a fluent percept of biological motion (Giese & Poggio, 2003; Lange & Lappe, 2006). In this regard it is important to note that, so far, research in autism has focused on bottom-up biological motion perception, with no direct investigation of top-down processing in autism.

To address the hypothesis of an autism-related deviance in top-down biological motion perception, we used a recently validated EEG frequency tagging paradigm (Cracco et al., 2022, 2023) to measure apparent biological motion perception in a sample of adults scoring low *vs* high on autism traits (Cracco et al., 2022). We replicated the findings of Cracco et al. (2022). That is, we found a stronger brain response for fluent sequences at the frequency that primarily captures movement processing (i.e. half cycle), and a stronger brain response for non-fluent sequences at the frequency that primarily captures body posture processing (i.e. full cycle). However, we found no support for our autism-related hypothesis, as we did not find a specific reduction in the effect of movement fluency on half cycle responses. Hence, we found no evidence for a diminished temporal integration of body postures into movements in adults scoring high on autism traits.

This study therefore provides first evidence that top-down processing of biological motion is not different in autism. However, more research is needed to confirm our results as this is the first study that directly tested this type of biological motion processing in autism. Moreover, the current study used a dimensional approach to autism by testing neurotypical participants who scored either high or low on autism traits. Hence, the findings should be replicated in a sample of individuals with an actual diagnosis of autism. To clarify, autism traits are continuously distributed throughout the whole population, with at the extreme end of the continuum a subpopulation of individuals who may receive a diagnosis of autism (Abu-Akel et al., 2019). Previous research has shown that applying a dimensional approach to autism in the neurotypical population can produce relevant insights about autism (Goris et al., 2017, 2021; Grinter et al., 2009; Nijhof et al., 2017; Robertson & Simmons, 2013; Stewart & Austin, 2009; Walter et al., 2009), Nevertheless, we cannot exclude the possibility that a diminished integration of body postures into movements is instead a categorical characteristic of autism that only emerges in individuals with a formal diagnosis of autism. Therefore, the current study needs to be followed-up by research in clinical samples.

While the results of our study did not support our hypothesis of autism-related diminished top-down processing of biological motion, we observed an interesting group effect. Specifically, we found that the group that scored high on autism traits showed a stronger fluency effect on both the half (fluent > non-fluent) and full cycle (non-fluent > fluent) response. This effect could be interpreted as enhanced temporal integration of body postures into movements in individuals that score relatively high on autism-traits – the opposite of what was expected. Importantly however, because the greater fluency effect was not specific to the half cycle response but was also there for the full cycle response, this interpretation would further imply that the high-scoring group showed enhanced body-posture perception as well. Another perhaps more parsimonious explanation is that there is a more general perceptual difference between individuals who score high *vs* low on autism traits. To clarify, the image sequences used in the current study could be perceived in two ways: as a sequence of body postures (repeating at full cycle) or as a sequence of movements (repeating at half cycle). Crucially, by manipulating fluency, we could make one percept more salient than the other (i.e., the half cycle percept in the fluent condition and the full cycle percept in the non-fluent condition). Therefore, a parsimonious explanation for the finding that both the half cycle and the full cycle responses were more sensitive to fluency in the high-scoring group is that this group was more influenced by perceptual saliency. While speculative, this potentially indicates that individuals scoring high on autism traits have a perceptual processing style that focuses more on a single percept, in this case the more salient one, whereas individuals scoring low on autism traits have a perceptual processing style in which the two possible percepts (fluent apparent movement or sequence of static body postures) are less clearly dissociated from each other.

This perceptual processing style found in our group scoring high on autism traits aligns well with the autism literature on perceptual stability and rigidity. Previous studies have shown that autism is related to a perceptual processing style that is ‘overly’ stable (Watanabe et al., 2019). Studies on multistable perception have for example used ambiguous figures (e.g. Necker cubes) that can result in two or more equally possible percepts and create spontaneous perception reversals (Long et al., 1951). Autism studies have found fewer such reversals in adults and children with autism compared to adults and children without autism (Kornmeier et al., 2017; Sobel et al., 2005), with some participants with autism reporting no reversals at all (Kornmeier et al., 2017). Similarly, Watanabe et al. (2019) also reported less reversals in adults with autism compared to adults without autism using bistable structure- from-motion stimuli (i.e. a field of moving dots that can be seen as a sphere that rotates clockwise or counterclockwise). Binocular rivalry studies, which show different images to each eye, also found less reversals in autism (Freyberg et al., 2015; Robertson et al., 2013). Furthermore, Allen and Chambers (2011) reported perceptual rigidity in the way adolescents with autism mentally represent ambiguous figures. More specifically, using a drawing task, they showed that drawings by adolescents with autism were less influenced by contextual biasing (a label of one of two percepts) compared to a matched control group. Our results may thus be explained by this broader phenomenon of perceptual stability known in autism, as opposed to an explanation specific to biological motion perception. On this view, people scoring high on autism traits perceive *either* fluent movement *or* a sequence of body postures, while people scoring low on autism are more like to perceive both percepts at the same time, or – alternatively – perceptually alternate between them over the time course of an experimental trial. The bias in the high-scoring group for the most salient percept as observed in our study, is also in line with a recent study that showed that the often observed altered local/global processing in autism may not be due to a stronger local processing over global processing bias but that differences in local/global processing in autism are driven by exaggerated salience effects (Baisa et al., 2020).

A limitation of the current study is that, in order to minimize the duration of the task, we did not include a non-human condition to control for animacy. However, a recent study by Cracco et al. (2023) that used the same paradigm with both a human (animate) and corkscrew (non-animate) condition found that movement processing did not depend on animacy. This is consistent with previous work on the bottom-up processing of biological motion (Jastorff et al., 2006 and with theoretical models of biological motion perception (Giese & Poggio, 2003; Lange & Lappe, 2006). Moreover, as outlined above, our findings are best explained by the broader phenomenon of perceptual stability which applies to the visual perception of both animate and inanimate stimuli. Nonetheless, future studies on biological motion perception in this area might want to add such a control condition.

To conclude, we did not find evidence for differences in top-down processing of biological motion in adults who score high *vs*. low on autism traits. Instead, we found that adults scoring high on autism traits have a more stable perceptual processing style/bias to process the more salient percept. This processing style is likely to be independent of biological motion perception and therefore points towards a more general perceptual difference that is not necessarily tied to social cognition. However, as this was the first study investigating top-down biological motion perception in the context of autism, further research with different paradigms and preferably with a sample of individuals that have a confirmed autism diagnosis, is warranted before firm conclusions can be drawn.

## Acknowledgements

The authors would like to thank all participants for their contribution as well as Anne de Groote and Hannah de Laet for their assistance with data collection, and their help with participant recruitment. DO was supported by the Special Research Fund of Ghent University (BOF18/DOC/348). GO received funding from the European Research Council (ERC) under the European Union’s Horizon 2020 research and innovation programme (grant agreement No. 864420 - Neurolive). EC was supported by a postdoctoral fellowship awarded by the Research Foundation Flanders (12U0322N).

## Conflict of interest

The authors declared that they had no conflict of interest with respect to their authorship or the publication of this article.

## Supplementary material

### Methods

#### Random condition check

As noted in the methods, section the random condition of Cracco et al. (2022) was administered as practice block in the current study. This control condition is supposed to only elicit a base rate response, as neither postures nor movements repeat predictably when images are presented at random. This was indeed the case in the current study, as we found a base rate response (*M* = 1.90) for the control condition, whilst the full and half cycle response was virtually zero (*M*_full_= 0.09; *M*_half_ = 0.04). These results are supported by the topographies (Supplementary Figure 1). Note that due to a procedural error we did not record EEG data of the practice block for three participants.

**Supplementary Figure 1.**
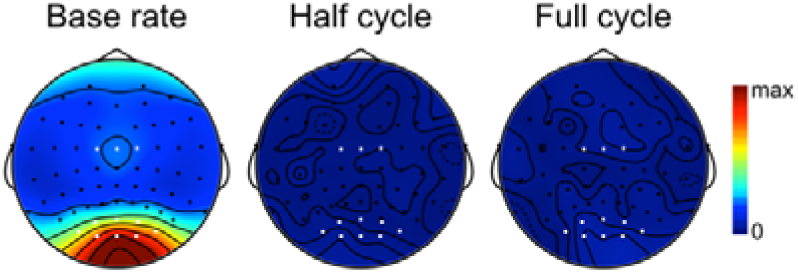
Topographies of the random condition for each frequency.

### Results

#### Full cycle

**Supplementary Table 1.** Means and standard deviations of the Condition × Region effect.

| Condition | Region | $M$ (SD) |
| --- | --- | --- |
| Non-fluent | Left posterior | 2.36 (1.02) |
|  | Middle posterior | 1.46 (0.76) |
|  | Right posterior | 2.51 (1.12) |
|  | Central | 0.33 (0.19) |
| Fluent | Left posterior | 1.79 (0.88) |
|  | Middle posterior | 1.20 (0.07) |
|  | Right posterior | 1.89 (0.79) |
|  | Central | 0.27 (0.20) |
| Fluency effect | Left posterior | -0.57 (0.75) |
|  | Middle posterior | -0.26 (0.55) |
|  | Right posterior | -0.62 (0.84) |
|  | Central | -0.06 (0.23) |

*Note.* Fluency effect = fluent – non-fluent

#### Half cycle

**Supplementary Table 2.** Means and standard deviations of the Condition × Region effect.

| Condition | Region | *M (SD)* |
| --- | --- | --- |
| Non-fluent | Left posterior | 2.10 (0.99) |
|  | Middle posterior | 2.47 (1.01) |
|  | Right posterior | 2.42 (1.08) |
|  | Central | 0.61 (0.32) |
| Fluent | Left posterior | 3.16 (1.21) |
|  | Middle posterior | 3.52 (1.25) |
|  | Right posterior | 3.64 (1.60) |
|  | Central | 1.12 (0.48) |
| Fluency effect | Left posterior | 1.06 (0.89) |
|  | Middle posterior | 1.04 (0.97) |
|  | Right posterior | 1.22 (1.03) |
|  | Central | 0.52 (0.38) |

*Note.* Fluency effect = fluent – non-fluent**Table 3***Means and standard deviations of the Region × Condition × Group effect*

**Supplementary Table 3.** Means and standard deviations of the Region × Condition × Group effect.

| Condition | Region | Group | *M (SD)* |
| --- | --- | --- | --- |
| Non-fluent | Left posterior | Low-scoring | 2.24 (1.15) |
|  |  | High-scoring | 1.97 (0.80) |
|  | Middle posterior | Low-scoring | 2.67 (1.08) |
|  |  | High-scoring | 2.28 (0.91) |
|  | Right posterior | Low-scoring | 2.59 (0.97) |
|  |  | High-scoring | 2.26 (1.17) |
|  | Central | Low-scoring | 0.60 (0.28) |
|  |  | High-scoring | 0.61 (0.36) |
| Fluent | Left posterior | Low-scoring | 3.16 (1.27) |
|  |  | High-scoring | 3.16 (1.17) |
|  | Middle posterior | Low-scoring | 3.46 (1.25) |
|  |  | High-scoring | 3.57 (1.27) |
|  | Right posterior | Low-scoring | 3.50 (1.42) |
|  |  | High-scoring | 3.78 (1.78) |
|  | Central | Low-scoring | 1.06 (0.40) |
|  |  | High-scoring | 1.19 (0.55) |
| Fluency effect | Left posterior | Low-scoring | 0.92 (0.96) |
|  |  | High-scoring | 1.20 (0.81) |
|  | Middle posterior | Low-scoring | 0.79 (0.91) |
|  |  | High-scoring | 1.29 (0.97) |
|  | Right posterior | Low-scoring | 0.91 (0.95) |
|  |  | High-scoring | 1.52 (1.04) |
|  | Central | Low-scoring | 0.46 (0.37) |
|  |  | High-scoring | 0.57 (0.38) |

*Note.* Fluency effect = fluent – non-fluent

## Footnotes

1 The late identification of faulty equipment led to this high number of participants with bad signal quality.

## Notes

### Competing Interest Statement

The authors have declared no competing interest.

